# Paired metagenomic and chemical evaluation of an aflatoxin contaminated dog kibble

**DOI:** 10.1101/2024.02.14.580305

**Authors:** Andrea Ottesen, Brandon Kocurek, Elizabeth Reed, Seth Commichaux, Mark Mammel, Padmini Ramachandran, Patrick McDermott, Brenna M. Flannery, Errol Strain

## Abstract

Identification of chemical toxins from complex or highly processed foods can present ‘needle in the haystack’ challenges for chemists. Metagenomic data can guide chemical toxicity evaluations with DNA-based description of the wholistic composition (bacterial, eukaryotic, protozoal, viral, and antimicrobial resistance) of any food suspected to harbor toxins, allergens, or pathogens. This approach can focus chemistry-based diagnostics, improve risk assessment, and address data gaps. There is increasing recognition that simultaneously co-occurring mycotoxins, either from single or multiple species, can impact dietary toxicity. Here we evaluate an aflatoxin contaminated kibble with known levels of specific mycotoxins and demonstrate that the abundance of DNA from putative aflatoxigenic *Aspergillus* spp. correlated with levels of aflatoxin quantified by Liquid Chromatography Mass Spectrometry (LCMS). Metagenomic data also identified an expansive range of co-occurring fungal taxa which may produce additional mycotoxins. Metagenomic data paired with chemical data provides a novel modality to address current data gaps pertaining to mycotoxin toxicity exposures, toxigenic fungal taxonomy, and mycotoxins of emerging concern.

## Background

The evaluation of new methodological approaches to identify microbiological and chemical hazards in human and animal foods is a central focus of FDA research. Advancing science to improve regulatory policy is a central dogma underpinning the Agency’s public health mission. Currently, genomic and metagenomic data are used for food safety applications such as source tracking pathogens[1-6], identification of adulterants, contaminants, toxins, allergens[7], and detection and prediction of antimicrobial resistance(AMR)[8]. A new frontier of utility for metagenomic data includes support for chemical toxicity evaluations through provision of wholistic compositional DNA-based profiles of macro (plant and animal) and micro (bacterial, fungal, viral, AMR) components of both unprocessed and highly processed human and animal foods.

Molecular tools have facilitated a paradigm shift in our understanding of the ecology of pathogenicity associated with human and animal foods[9]. A similar trend in mycotoxin risk assessment has evolved to consider simultaneous co-exposure to diverse mycotoxins[10]. Mycotoxins are secondary metabolites produced by fungi. There are hundreds of different moieties produced by single and/or multiple species. It is rare for a single mycotoxin to exist in any crop or food due to the complex biodiversity of agricultural ecologies[11]. Reported fungal species for corn alone include-*Fusarium proliferatum, Trichoderma gamsii, T. longibrachiatum, Penicillium oxalicum, P*.*aurantiogriseum, P. polonicum, Bipolaris zeicola, Sarocladium zeae, Chaetomium murorum, Botrytrichum murorum, Cladosporium cladosporioides, C. sphaerospermum, Aspergillus niger, A. flavus, Alternaria alternata, and Rhizopus microsporus*[12] - at least half of which are known to produce toxins of significance to human and animal health.

Risk assessment of mycotoxins in foods to date has primarily focused on a small number of important toxins with critical adverse effects, without considering the significance of co-exposure to multiple compounds[10] despite co-occurrence of diverse mycotoxins being more common than exposure to single mycotoxins. Recent studies of mycotoxin co-occurrence in animal feed in Europe have found that 75 to 100% contain more than one mycotoxin[13].

Exposure to a single toxin may exert multiple adverse effects and exposure to a mixture of structurally related or unrelated mycotoxins may also result in a combination of adverse effects[10]. Synchronous exposure to diverse toxins in food and feed may impact states of health and disease in ways we have yet to fully characterize.

*Aspergillus flavus* and A. *parasiticus* are two of the most traditionally recognized aflatoxigenic species in pre- and post-harvest commodities but there are numerous other aflatoxin producing species of *Aspergillus*, and it has even been proposed that species of *Fusarium, Penicillium, Claviceps*, and *Alternaria* may produce aflatoxins[14]. The International Agency for Research on Cancer (IARC) has classified aflatoxins as the most potent natural carcinogens known to humankind and they are estimated to contaminate 25% of crops worldwide. Many ingredients used for animal food, such as corn, wheat, and rice are some of the most susceptible to contamination by mycotoxins.

Whole genome sequencing (WGS) and metagenomic sequencing (MGS) have been used by state and federal agencies for almost two decades to describe how pathogens and toxins become associated with human and animal foods[1, 9, 15-19] but a new frontier of integrated chemical and metagenomic analyses is on the horizon for modernized stewardship of human and animal foods. Here we use MGS data to demonstrate that the taxonomic abundance of putative aflatoxigenic fungal species correlates with the levels of aflatoxin quantified by chemical methods (LCMS). Two levels of aflatoxin contaminated kibble quantified by LCMS were evaluated, one at 15ppb, and one at 522 ppb. (The 15 ppb kibble is below the FDA action level for aflatoxins of 20 ppb in pet food)[20]. Additionally we demonstrate that a wide range of species which may produce additional mycotoxins can be identified by the same data, to better address current data gaps and modernize risk assessment.

## Results

### Kibble ingredients identified by metagenomic sequencing

To describe composition and relative abundance of ingredients, metagenomic data were created for control, low-concentration (15 ppb), and high-concentration (522 ppb) aflatoxin contaminated kibble. The relative abundance of macro ingredients (annotated by mitochondrial DNA and occurring at greater than 1% of normalized data) included: Zea (corn), *Gallus* (chicken), *Triticum* (wheat), Soya, (soybean), Bos (cow, ox, bull, yak, cattle) and yeast (Figure 1A). Further refinement of species annotations described; *Z. mays, G. gallus* and *G. gallus ssp. spadiceus, T. aestivum, B. taurus*, several species of *Saccharomyces* (*cerevisiae, pastorianus* and *pastorianus Weihenstephan*) and *Fusarium verticillioides*. (Figure 1B).

**Figure 1.**
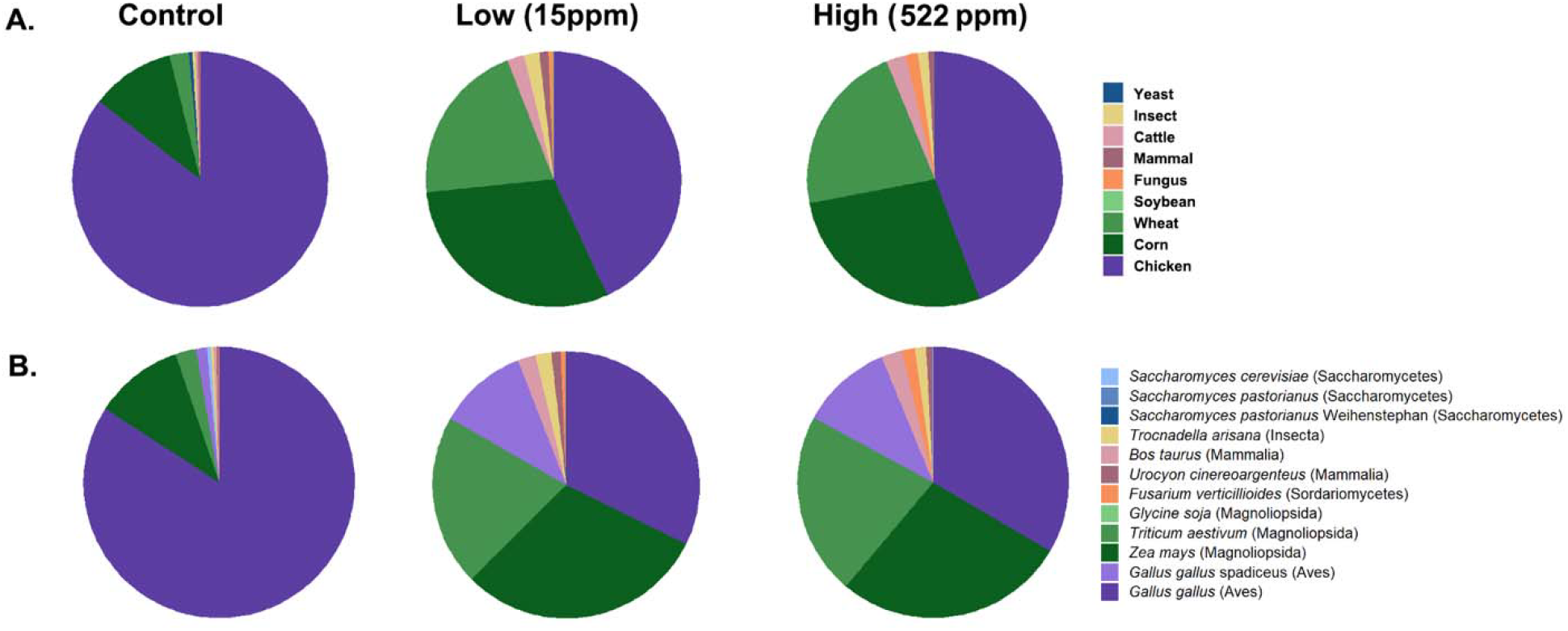
DNA based identification of eukaryotic components of control and low and high level aflatoxin contaminated kibble. 1A) DNA based identification of eukaryotic components of control, low, and high level aflatoxin contaminated kibble. 1B) Higher resolution species and serovar level identification of constituents in kibble. Ingredients of kibble were annotated using a database of mitochondrial sequences with a k-mer based detection approach.

### *Aspergillus* species in aflatoxin contaminated kibble

Evaluation of the metagenomic data from control, low, and high level aflatoxin containing kibble (quantified by LCMS), correlated with relative abundance of DNA of *Aspergillus* species, i.e.; the low level aflatoxin contaminated kibble had a low abundance of DNA from *Aspergillus* species and the high level kibble had a higher relative abundance of Aspergillus species. Almost no *Aspergillus* DNA was detected in controls (Figure 2). The *Aspergillus* species identified in control kibble was predominantly A. glaucus. Figure 2 shows the top twenty most abundant *Aspergillus species* identified in control, low (15 ppb), and high (522 ppb) levels of aflatoxin contaminated dog kibble. There was an extensive taxonomic range of *Aspergillus* species identified in the kibble. Production of aflatoxin may have been associated with a single species or potentially multiple species. *Aspergillus oryzae, flavus*, and *phoenicis* were the most abundant species observed in high level (522 ppb) aflatoxin contaminated kibble. *Aspergillus flavus* and A. parasiticus are two of the most traditionally recognized producers of aflatoxin in pre- and post-harvest commodities[14]. Figure 2B shows the relative abundance of A. *parasiticus, novoparasiticus*, and *flavus* without other *Aspergillus* species. These three species likely played a key role in aflatoxin production in the kibble as A. *oryzae* and A. *phoenicis* are not known to produce aflatoxins[21, 22].

**Figure 2.**
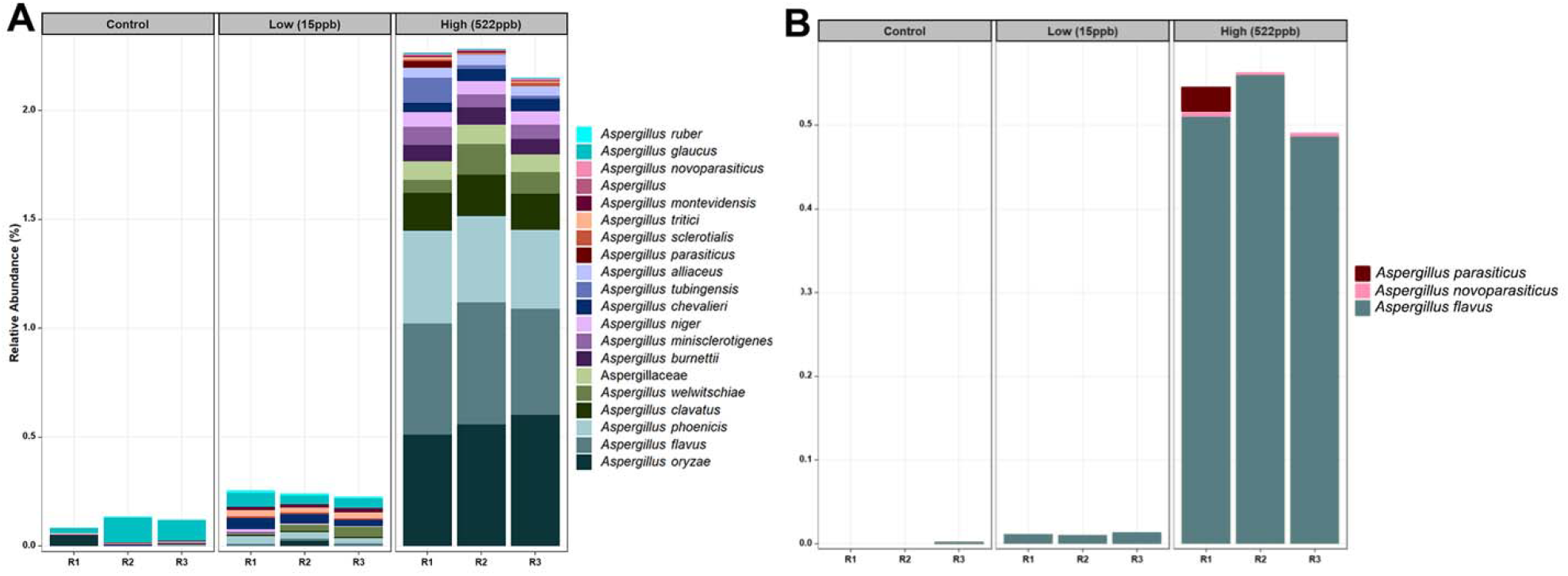
Relative abundance of the top twenty *Aspergillus* species and relative abundance of *Aspergillus* flavus, parasiticus, and *novoparasiticus* in control, low and high concentration kibble. 2A) Relative Abundance (Percentage) of the top twenty *Aspergillus* species and 2B) relative abundance of *Aspergillus flavus, parasiticus*, and *novoparasiticus* in control, low and high concentration kibble annotated using an in house FDA Fungal Database V1.2.

### Additional mycotoxin detection and associated species

While aflatoxin B1 was the primary focus of the chemical evaluation of the kibble due to its acute toxicity, concentrations of fumonisins (B1,B2,B3), and ochratoxin A were also detected by LCMS (Table 1).

**Table 1.**
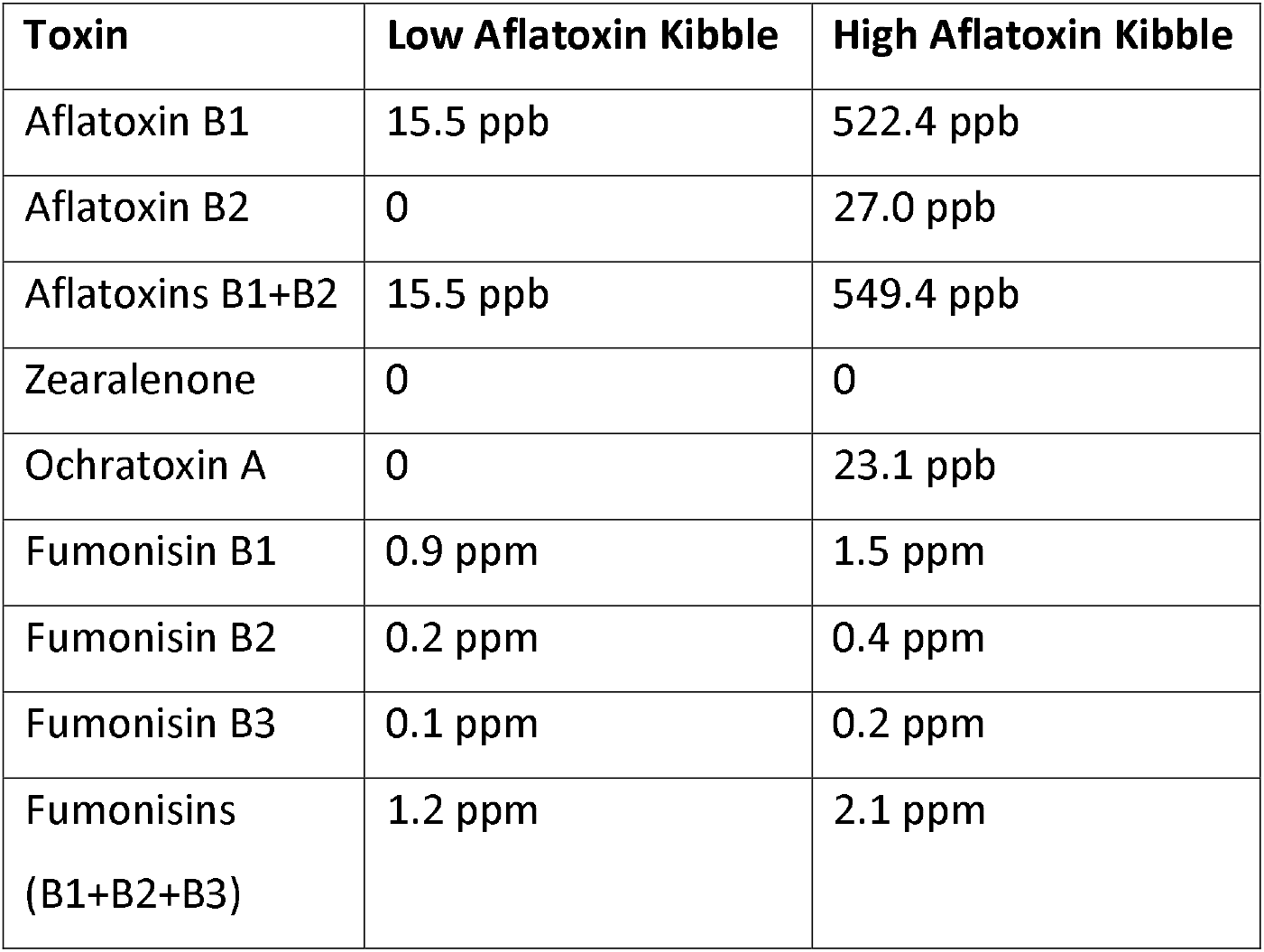
Chemical concentrations of mycotoxins in low and high levels of aflatoxin contaminated dog kibble.

| Toxin | Low Aflatoxin Kibble | High Aflatoxin Kibble |
| --- | --- | --- |
| Aflatoxin B1 | 15.5 ppb | 522.4 ppb |
| Aflatoxin B2 | 0 | 27.0 ppb |
| Aflatoxins B1+B2 | 15.5 ppb | 549.4 ppb |
| Zearalenone | 0 | 0 |
| Ochratoxin A | 0 | 23.1 ppb |
| Fumonisin B1 | 0.9 ppm | 1.5 ppm |
| Fumonisin B2 | 0.2 ppm | 0.4 ppm |
| Fumonisin B3 | 0.1 ppm | 0.2 ppm |
| Fumonisin<br>(B1+B2+B3) | 1.2 ppm | 2.1 ppm |

*Fusarium verticillioides* is reported to produce fumonisins and zearalenone, and was observed in both low and high samples of kibble but not in controls (Figure 3A). *Additional Fusarium* species including *F. annulatum, proliferatum, dlaminii, siculi, irregulare, pilosicola*, and *globosum* were primarily associated with contaminated kibble and were not observed in controls (Figure 3A). Deoxynivalenol is also produced by Fusarium species, but not measured in this study. Because chemical data was not collected for control kibble, it is difficult to speculate how the abundance of *Fusarium verticillioides* in both low and high kibble may have impacted the chemical profiles of the kibble.

**Figure 3.**
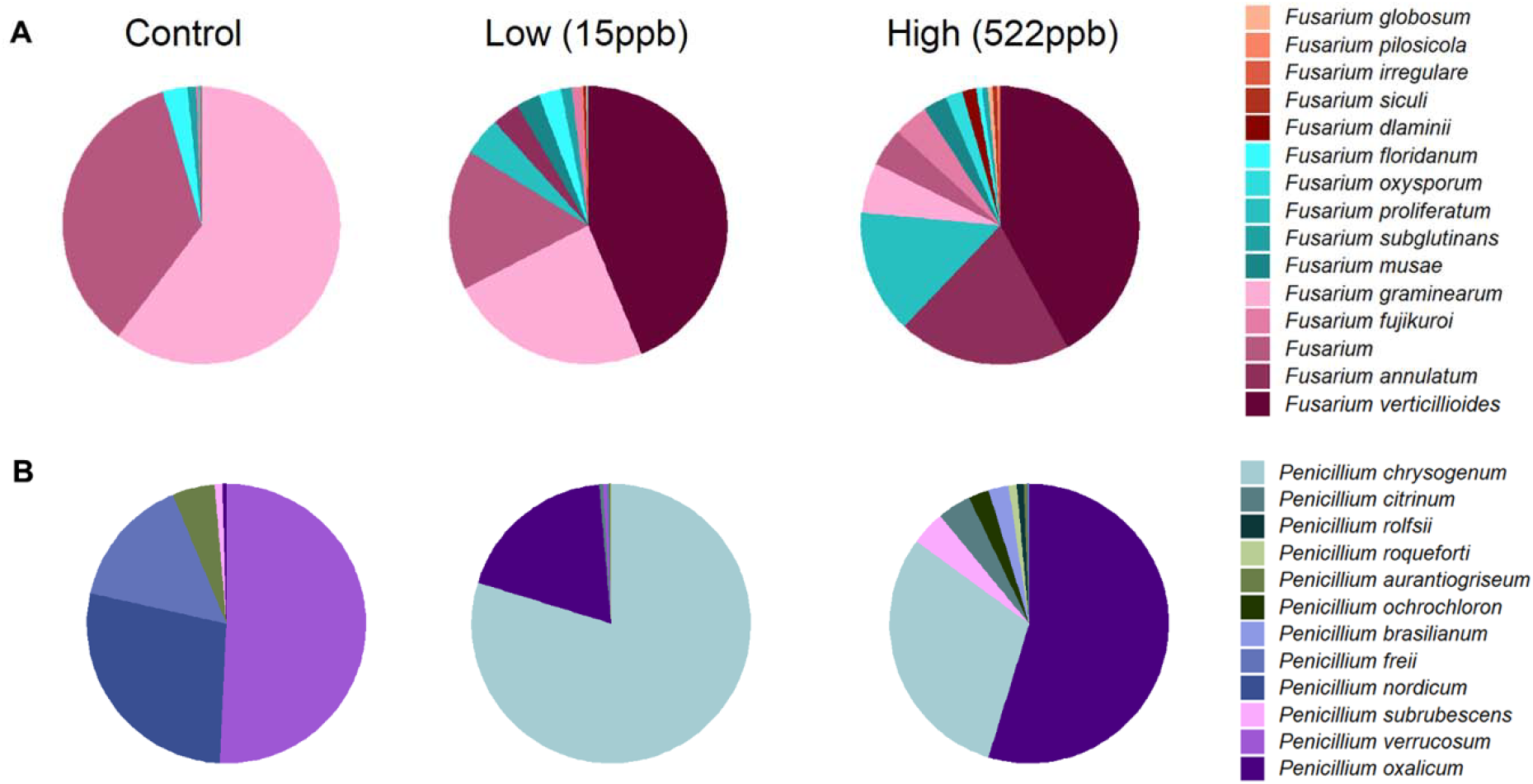
Relative abundance of *Fusarium* and *Penicillium* species in control, low, and high levels of aflatoxin contaminated kibble. Figure 3A shows the average relative abundance of *Fusarium* species from three replicates of each kibble type, (control, low, and high level aflatoxin contamination) annotated using an in house FDA bacterial database developed for metagenomic data. Figure 3B shows the average relative abundance of *Penicillium* species in each kibble type. Contaminated kibble had a distinctive incidence of *Fusarium verticillioides, Penicillium oxalicum* and *P. chrysogenum*, not seen in control kibble.

*Penicillium spp*. are reported to produce Ochratoxin A and patulin[23]. *Penicillium oxalicum, subrubescens, brasilianum, ochrochloron, roqueforti*, and *citrinum* were primarily observed in contaminated samples whereas *P. verrucosum*, nordicum and *freii* were the most abundant species observed in controls (Figure 3B). It is possible that *P. oxalicum* or *P. subrescens* played a role in the observed ochratoxin levels (23 ppb) in ‘high’ samples. However, *P. oxalicum* was also observed in low level samples for which no ochratoxin was quantified, so perhaps other genera differentially enriched in high level samples were responsible for the ochratoxin observed by LCMS (Table 1).

### Linear discriminant analysis of fungal species in low and high level contaminated kibble

Using linear discriminant analysis (LDA) from the COSMOSID analysis pipeline (CosmosID Metagenomics Cloud, app.cosmosid.com, CosmosID Inc., www.cosmosid.com) to compare differential abundance of fungal species in low, control, and high level aflatoxin contaminated kibble, we observed that high level kibble contained a greater abundance of *Aspergillus* species than low level (Figure 4). Interestingly, LDA highlights that control kibble had significantly more *Saccharomyces* species than the high level kibble. It is common practice to add yeast to ingredients that may contain levels of aflatoxin. Yeast species have been shown to bind aflatoxin B and thus reduce the toxic impact of consumption of aflatoxin contaminated food[24]. This approach has been shown to provide a protective effect to broiler chickens consuming aflatoxin B1 contaminated food[25]. Yeast was listed as an ingredient in all dog kibble.

**Figure 4.**
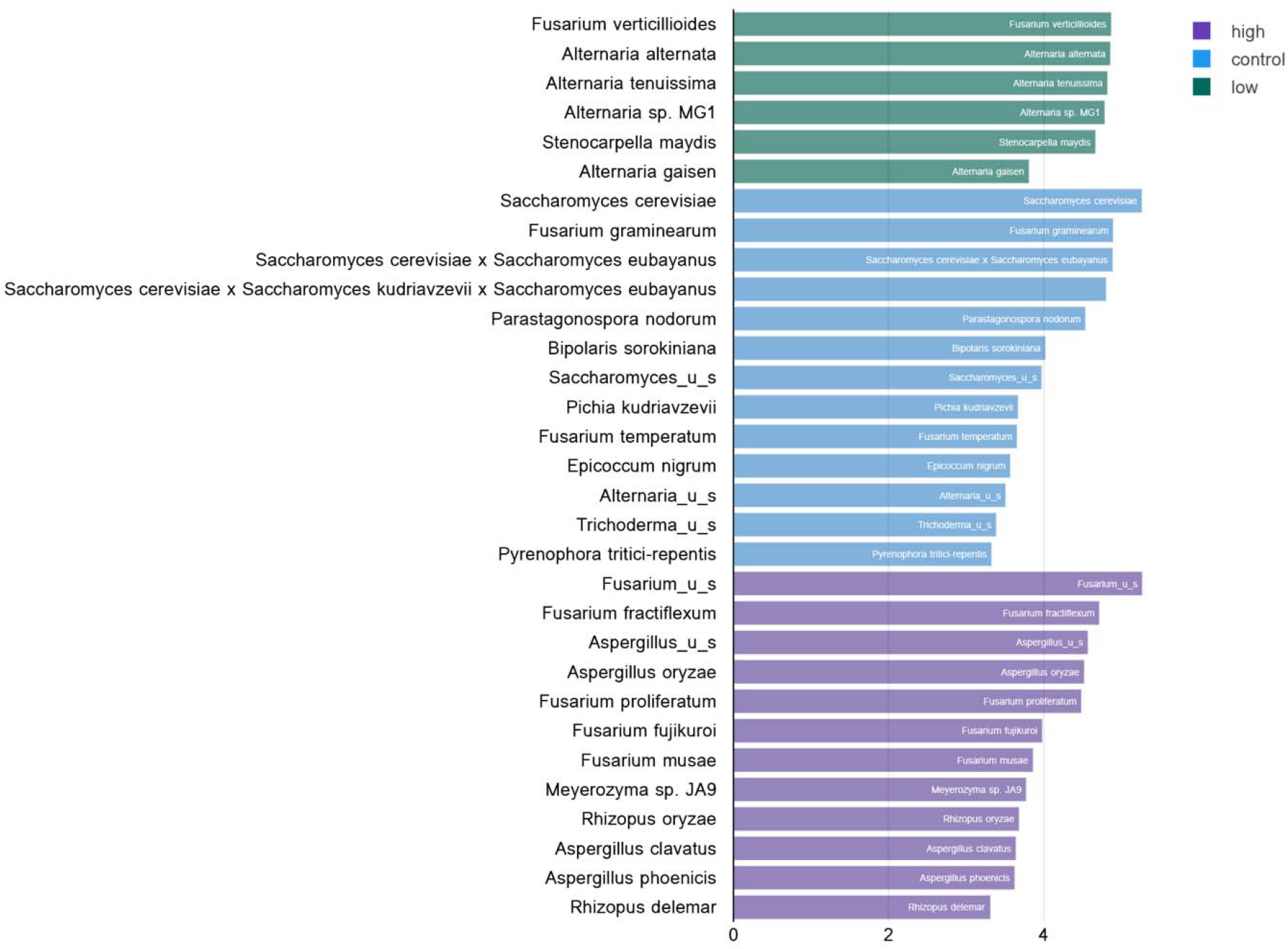
Linear discriminate analysis (LDA) of significant (p<0.05) differential abundance of fungal species in low, control, and high level aflatoxin contaminated dog kibble. Using the COSOMOSID annotation pipeline(CosmosID Metagenomics Cloud, app.cosmosid.com, CosmosID Inc., www.cosmosid.com) taxa that were significantly enriched in each type of kibble (control, low, high) were described. The LDA threshold spanned 2.9 - 5.27 with a p value range of 0.022-0.035.

Additionally, a greater relative abundance of *Alternaria* species were observed in low level aflatoxin contaminated kibble. While *Alternaria* species were observed in metagenomic data, *Alternaria*-associated toxins (altenuene, alternariol, alternariol monomethylether, tentoxin, and tenuazonic acid) were not measured in this study. Figure 5 provides a breakdown of the species of *Alternaria* across each kibble type.

**Figure 5.**
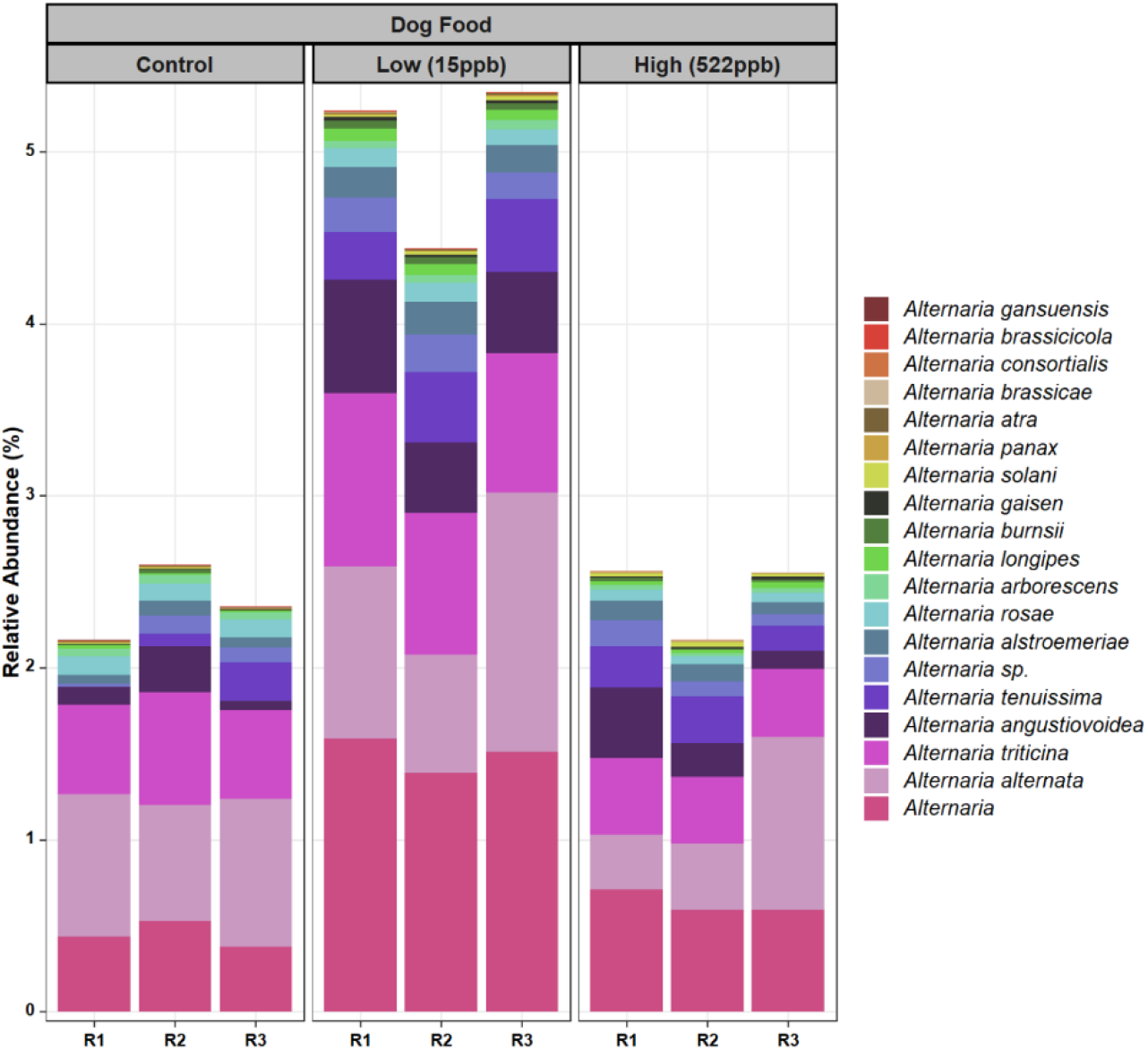
Species of *Alternaria* identified in control, low, and high level aflatoxin contaminated kibble. Species of *Alternaria* observed in the control, low, and high level aflatoxin contaminated kibble are shown for each of three replicates. For the most part, similar species were observed across all kibble types, however a greater relative abundance was observed in low level samples.

For *Alternaria*, there lacked a clear differentiation of species composition from control to contaminated kibble as was observed for genera like *Aspergillus, Penicillium*, and *Fusarium*. Fingerprints of differential abundance of fungal families that play important roles in food safety (Figure 6) could potentially be used to quickly detect lots of feed or ingredients that may be at risk of dangerous levels of aflatoxins or other significant mycotoxins or even to identify failed critical controls. When *Saccharomyces* species (used to bind aflatoxin) have been depleted, there is likely a greater risk of toxicity exposure. Targeted assays using Polymerase Chain Reaction (PCR) based methods have provided effective identification of species known to produce mycotoxins [26-28]. However, PCR based approaches cannot describe the wholistic composition of food, cannot identify novel taxa, and cannot describe multi-taxa relative abundances that may prove critical to the safety of the food.

**Figure 6.**
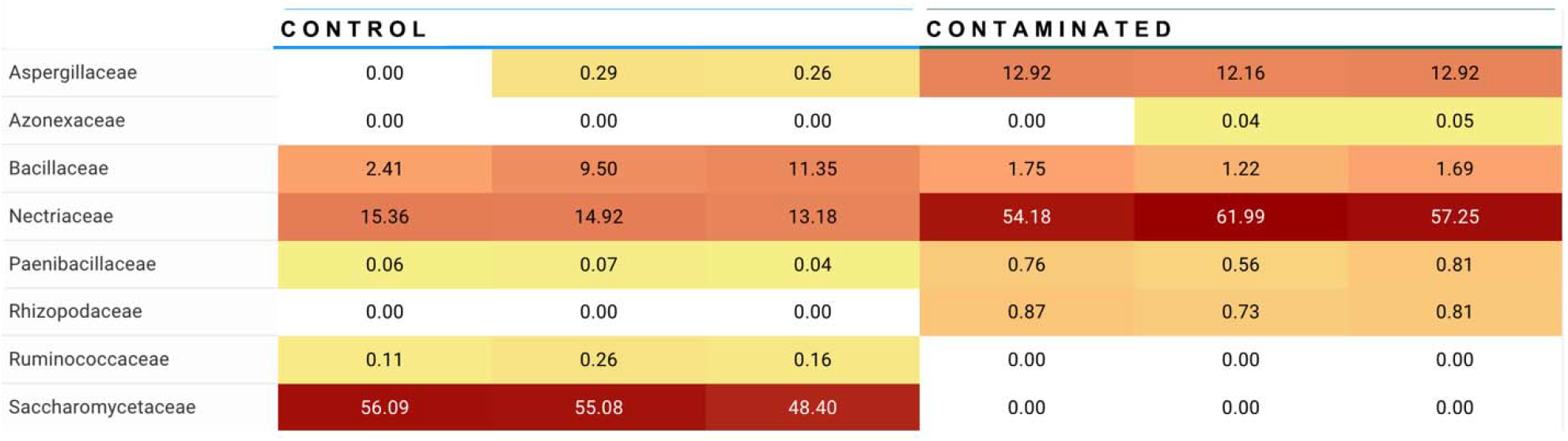
Summary of differential abundance at fungal family level. Families Apergillaceae (containing *Aspergillus* genera) and Nectriaceae (containing *Fusarium* genera) were observed at significantly higher abundance in contaminated kibble (high level aflatoxin) than in control kibble, and Saccharomycetaceae (containing *Saccharomyces* genera i.e., yeast) are at significantly higher abundance in control kibble contrasted to contaminated kibble.

### Identification of functional genes in mycotoxin pathways

Beyond identification of taxonomic structure of kibble, metagenomic data also described functional genes involved in mycotoxin production and thus provided further confirmation of mycotoxin pathways such as; deoxynivalenol, nivalenol, and aflatoxin. Not surprisingly, the most numerous hits to genes involved in the aflatoxin production pathway were seen in the high level aflatoxin contaminated samples (Table 2). The doublet and singleton hits to nivalenol (NIV) and deoxynivalenol (DON) in control samples may be indicative of low levels of NIV or DON in these samples. NIV is a mycotoxin of the trichothecene group produced by *Fusarium* species. DON, also referred to as vomitoxin, is also produced by *Fusarium species*, notably; *graminearum*[29], which was seen in high abundance in control samples (Figure 3).

**Table 2.**
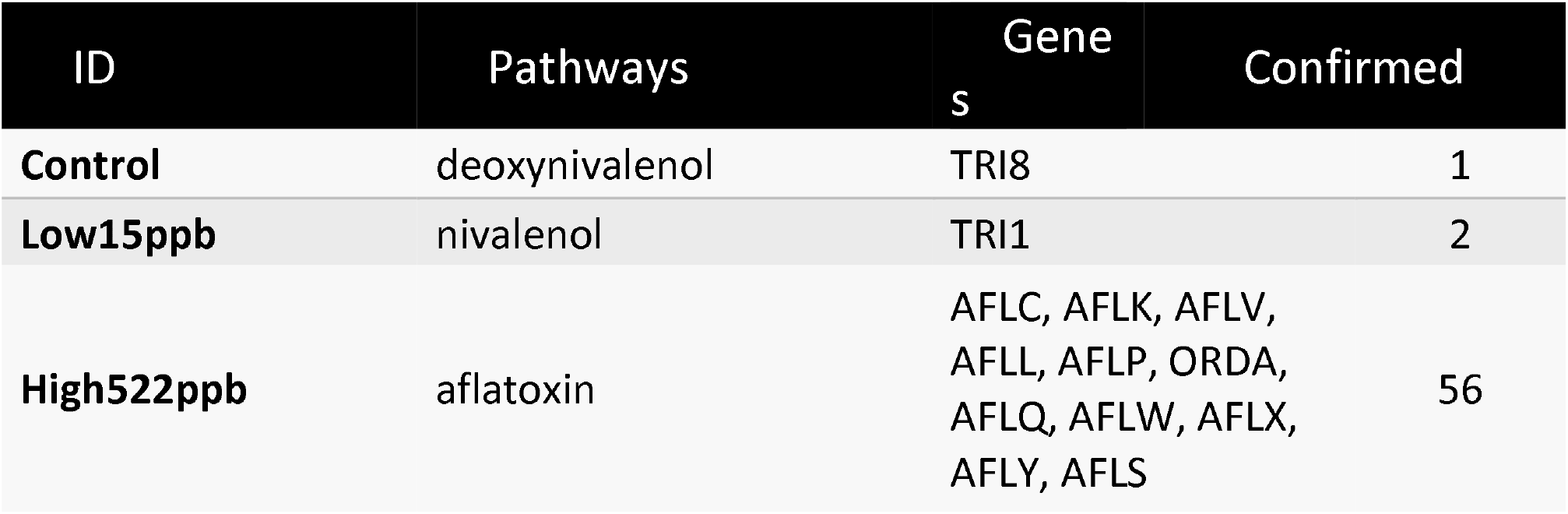
Hits to genes in deoxynivalenol, nivalenol, and aflatoxin pathways in metagenomic data of kibble. The number of hits to genes in mycotoxin production pathways for deoxynivalenol, nivalenol, and aflatoxin is show in the “confirmed” column. Hits were blasted to NCBI to confirm gene identification.

### Additional interesting observations

*Stenocarpella maydis*, another potential toxin producer, was observed in contaminated kibble samples but not in controls. *Stenocarpella maydis* is a fungal pathogen of corn and its toxic metabolites include diplodiatoxin, chaetoglobosins K and L, and (all-E)-trideca-4,6,10,12-tetraene-2,8-diol [30, 31]. Synonyms for *S. maydis* include: *Diplodia zeae, Diplodia maydis, Sphaeria maydis, S. zeae, Macrodiplodia zeae*, and *Dothiora zeae* which have been linked to diplodia toxicity (diplodiosis). Diplodiosis is characterized by muscle tremors, incoordination, ataxic hindquarters, paralysis, and death of cattle, sheep, rats, and ducklings, with reports of cattle mortalities dating back to 1919 [32]. Additional taxa that were unique to high level aflatoxin contaminated samples included *Rhizopus delemar* (synonym *R. oryzae*) which has been associated with mucormycosis leading to high fatality rates in immunocompromised individuals and people with diabetes mellitus[33]. There are also reports of mucormycosis contributing to fatalities in dogs[34].

### Bacterial species in metagenomes of dog kibble

Bacterial profiles for controls and aflatoxin contaminated dog kibble are shown in Figure 7. Three replicates of control, low, and high level aflatoxin contaminated dog kibble were averaged and taxa occurring at greater than 3% of total data were graphed to summarize bacterial features of each kibble. *Enterobacter, Serratia, and Kosakonia genera were observed* in toxin contaminated kibble and not in control kibble.

**Figure 7.**
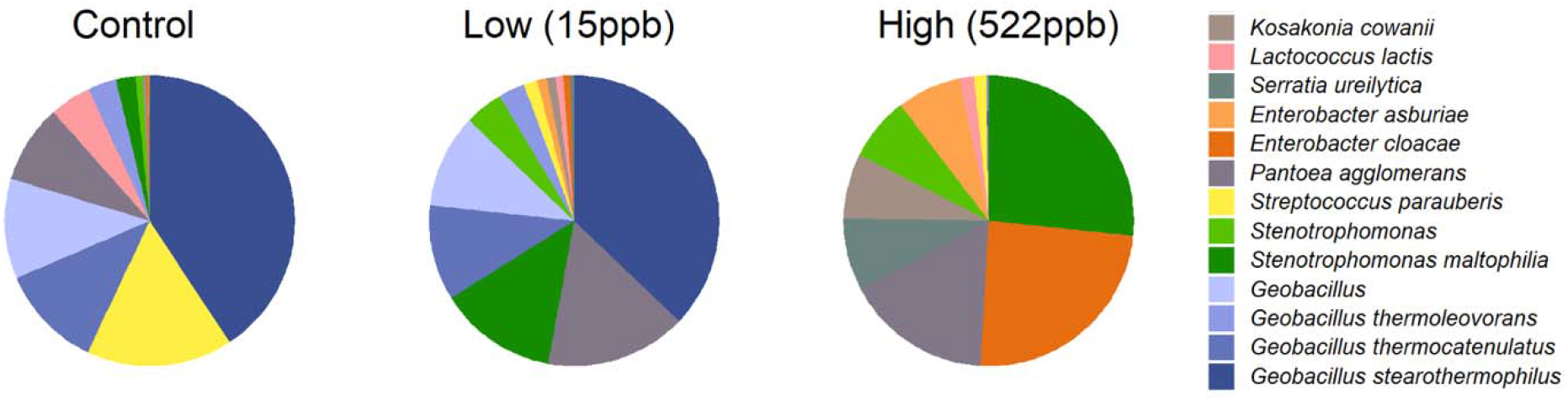
Bacterial Species in Control, Low and High Level Aflatoxin Contaminated Kibble. Bacterial taxa from control, low, and high level-aflatoxin contaminated dog kibble are shown here. The high level kibble had a strong incidence of *Enterobacter cloaceae. Stenotrophomonas* was also observed in kibble and has been described as an emerging (at times multi-drug resistant) global pathogen.

## Discussion

Aflatoxin outbreaks related to pet food have caused numerous companion animal deaths in the United States[35].[35]. A primary goal of the Center of Veterinary Medicine (CVM) is to protect human and animal health by assuring the safety of animal feed and companion animal food. Metagenomic data can contribute to this goal by providing a method to generate a comprehensive summary of all ingredients in any animal food, including high resolution species and subspecies level identification of animal, plant, fungal, and bacterial constituents of the food. This type of information paired with chemical toxicity profiles has the potential to address data gaps in our understanding of mycotoxin contamination.

Here we demonstrate that metagenomic data from dog kibble with low and high levels of aflatoxin contamination had a correlative abundance of DNA from putative toxin producers *Aspergillus flavus, A. parasiticus*, and *A. novo-parasiticus*. This observation supports the hypothesis that these species played a key role in the aflatoxin contamination of the kibble. Additional correlative chemical and molecular benchmarking may be used for future safeguarding measures and potentially to accurately predict contamination levels of specific toxins. MGS used in concert with chemical profiling will not only be useful to predict contamination by known aflatoxigenic species but also to identify new species of mycotoxigenic fungi in feed in food.

Additionally, MGS data can identify fungal species in food and feed regionally and temporally, which may be useful to identify shifts in fungal contaminants occurring in response to global climate changes[22]. Furthermore, this approach coupled with focused enrichment of *Aspergillus species* could potentially identify *Aspergillus strains* that may be resistant to azole antibiotics which could be significant to *Aspergillus* pulmonary infections.

FDA currently monitors for aflatoxins, fumonisins, vomitoxin, zearalenone and ochratoxin A in human and animal food using targeted methods that detect these chemicals[36, 37]. A multi-mycotoxin surveillance approach has recently been initiated by FDA for infant and toddler foods[38]. This approach could benefit greatly from paired metagenomic and chemical data. MGS of food is a non-targeted method that provides a comprehensive characterization of all organisms present in any sample, including mycotoxigenic fungi that produce mycotoxins not currently monitored by FDA. Therefore, MGS results could signal the potential for additional mycotoxin testing, especially if a food has been associated with illness. Indeed, results from the dog kibble described multiple *Penicillium* species, which can produce toxins such as brevianamid A, citreoviridin, citrinin, cyclopiazonic acid, fumitremorgin B, griseofulvin, luteoskyrin, *penicillic* acid, penitrem A, PR-toxin, roquefortine, rugulosin, verrucosidin, verruculogen, viridicarumtoxin and xanthomegnin[39]. Furthermore, results demonstrated presence and abundance of toxin producing fungi *Stenocarpella maydis, Rhizopus delemar and R. oryzae* in contaminated kibble and not in control kibble. Follow-up chemical analysis could potentially determine the presence of toxins associated with those fungi.

Use of ever-improving metagenomic tools to evaluate human and animal food can inform on a myriad of elements – from the identification of pathogens and toxins in agricultural growing environments to adulterants and contaminants introduced at points along manufacturing and distribution chains. Such data can describe the total composition (genus, species, subspecies and even serovars and varieties) of plants, animals, bacteria, fungi, viruses, plasmids, and associated antimicrobial resistance. The paired chemical and MGS methodology provides valuable information to underpin modernized risk assessment for human and animal foods and to support chemical based multi-mycotoxin assessment with correlative taxonomic profiling.

## Methods

### Kibble

Control, low, and high aflatoxin containing kibble samples were all from the same brand of dog food. Kibble ingredients listed chicken by-product meal, corn, wheat, meat meal, rice bran, chicken fat, dried beet pulp, whitefish meal, flaxseed, salt, potassium chloride, choline chloride, vitamins and minerals. The label described a composition of at least 26% protein, 15% fat with maximum fiber at 6%,moisture at 10%, and 3645 kcal/kg.

### Metagenome preparation

Three 2.5 g portions of each level of aflatoxin contaminated kibble and control were ground to a fine powder in a Qiagen Tissue Lyser II at 30 Hz (1800 oscillations per minute) for 1.3 minutes per sample. Replicates of 100 mg of powder from the pooled mixture (7.5 gram) were used for DNA extraction. DNA extraction was performed using the Zymo High Molecular Weight DNA extraction kit according to the manufacturer’s specifications including an extra extraction lysing step using PBS and lysozyme. Libraries of DNA were created using the Illumina DNA Library Prep Kit according to the manufacturer’s specified protocols and sequencing was performed on a NextSeq 2000 using a high throughput kit. An additional NextSeq 2000 high throughput sequencing run was performed with only 2 replicates of low, medium and high level samples to achieve a sequencing depth of between 100 and 250 million reads per replicate to evaluate how increased read depth impacted incidence of key species.

### Bioinformatic Analyses

Sequence data were analyzed using in house FDA pipelines and databases and also with the Cosmos ID cloud based applications (CosmosID Metagenomics Cloud, app.cosmosid.com, CosmosID Inc., www.cosmosid.com) with Fungal Database Version 1.2[40]. Sequence data annotations were visualized using graphs created by the R Tidyverse package (https://cran.r-project.org/web/packages/tidyverse/index.html).

### Mycotoxin pathway identification

Metagenomic reads were mapped with Diamond (v2.0.5)[41] BLASTX (≥95% identity and those ≥90% read coverage) to a database of genes involved in mycotoxin biosynthetic pathways: aflatoxin, deoxynivalenol, nivalenol, ochratoxin, patulin, sterigmatocystin, T2_toxin, tenuazonic_acid. Genes were identified and downloaded from the MetaCyc database[42] (https://metacyc.org/). Identity of reads aligning to mycotoxin genes was confirmed by BLASTX aligning them to the NCBI nr database online.

### Mycotoxin evaluation

HPLC aflatoxin evaluation was conducted according to standard operating procedures of the Plant Industries Division of the Missouri Department of Agriculture which uses the AOAC Official method 2005.08: Aflatoxins in Corn, Raw Peanuts, and Peanut Butter (Liquid chromatography with Post-Column Photochemical Derivatization). An additional LCMS Multiple Mycotoxin evaluation was conducted by a 3^rd^ party Feed Evaluation Lab (Cumberland Valley Analytical Services, Zullinger, PA). Mycotoxins evaluated included aflatoxin, fumonisin, ochratoxin A and zearalenone in control and contaminated kibble.

## Acknowledgements

We would like to thank Stan Cook, Madison Fink, Quintin Muenks, and Mary Koestner of the Missouri Department of Agriculture - Plant Industries, Bureau of Feed and Seed for provision of research samples of kibble.

## Data Availability

The data presented here have been deposited at the National Center for Biological Information (NCBI) under the Companion Animal Food Metagenome Bioproject (BioProject <u>PRJNA1062328</u>) with accession numbers <u>SAMN39295889</u> through <u>SAMN39295903</u>. The data was deposited at NCBI using the Genomic Standards Consortium (GSC) Minimum Information about any Sequence (MIxS) compliant MIMS food-animal and animal feed package (version 6).

